# Multiple Mutualist Effects generate synergistic selection and strengthen fitness alignment in a tripartite interaction between legumes, rhizobia, and mycorrhizal fungi

**DOI:** 10.1101/2021.01.26.428300

**Authors:** Michelle E. Afkhami, Maren L. Friesen, John R. Stinchcombe

**Affiliations:** Department of Biology, University of Miami, 1301 Memorial Dr. #215, Coral Gables, FL 33146; Department of Plant Pathology, Department of Crop and Soil Sciences, Washington State University, 321 Johnson Hall, Pullman WA 99164 USA; Department of Ecology and Evolutionary Biology, University of Toronto, 25 Willcocks St., Toronto, ON, Canada M5S 3B2

**Author notes:** **Statement of Authorship:** All authors conceived the overall project, contributed to the development of experimental plans, and to the manuscript draft. MEA conducted the experiment and collected the data with advice from JRS. MEA and JRS analyzed the data; funding to MEA and JRS supported experimental work, with funding to all authors supporting manuscript writing and analysis.

**Keywords:** multiple mutualist effects, rhizobia, mycorrhizal fungi, Medicago, nonadditive, genotypic selection analysis, symbiosis, fitness

## Abstract

Nearly all organisms interact with multiple mutualists, and complementarity within these complex interactions can result in synergistic fitness effects. However, it remains largely untested how multiple mutualists impact eco-evolutionary dynamics. We tested how multiple microbial mutualists-- N-fixing bacteria and mycorrrhizal fungi-- affected selection and heritability in their shared host plant (*Medicago truncatula*), as well as fitness alignment between partners. Our results demonstrate for the first time that multispecies mutualisms synergistically affect selection and heritability of host traits and enhance fitness alignment between mutualists. Specifically, we found that multiple mutualists doubled the strength of selection on a plant architectural trait, resulted in 2-3-fold higher heritability of reproductive success, and more than doubled the strength of fitness alignment between N-fixing bacteria and plants. Taken together, these findings show that synergism generated by multiple mutualisms extends to key components of microevolutionary change, and emphasizes the importance of multiple mutualist effects in understanding evolutionary trajectories.

## Introduction

Mutualisms - associations in which all participants benefit - dramatically affect the interacting species as well as the dynamics of their populations, communities, and ecosystems (Bronstein 2015; Chomicki *et al.* 2019; David *et al.* 2019). Most organisms participate in many mutualisms throughout their lifetime or even simultaneously (McKeon *et al.* 2012; Afkhami *et al.* 2014). For instance, plants may associate with pollinators, seed dispersers, ant defenders, foliar endophytes, and rhizosphere nutritional symbionts. While traditionally the study of these beneficial interactions has been pairwise, recently there has been a growing body of empirical and theoretical research exploring the ecological consequences of ‘Multiple Mutualist Effects’ (MMEs) (Stanton 2003; McKeon *et al.* 2012; Afkhami *et al.* 2014; Ossler *et al.* 2015; Keller *et al.* 2018). MMEs are common, and often generate nonadditive effects on performance or fitness of shared hosts (Afkhami *et al.* 2014) as well as on the molecular phenotypes underpinning these effects (Afkhami & Stinchcombe 2016; Palakurty *et al.* 2018). In particular, MMEs, unlike effects of multiple predators (or competitors), can synergistically increase fitness of shared hosts if each partner species provides distinct and complementary rewards (e.g., rhizobia and mycorrhizal fungi providing nitrogen and phosphorus, respectively, to a host plant; Afkhami *et al.* 2020). Although nonadditive effects can alter ecological and evolutionary dynamics of species in ways that cannot be predicted from studies of pairwise interactions, our understanding of how they do so remains limited. Here we examine how multiple microbial mutualists-- N-fixing bacteria and mycorrrhizal fungi--affect trait selection and heritability in their shared host plant (*Medicago truncatula*), as well as fitness alignment between partners.

One important way that MMEs could have significant consequences for the evolutionary trajectories of participating organisms is through nonadditive selection (TerHorst *et al.* 2015; Bolstad 2017; terHorst *et al.* 2017). For evolution to be driven solely by pairwise effects, traits involved in one interaction must evolve independently of other mutualistic partners, meaning that selection on those traits would be unaffected by multiple mutualists (Iwao & Rausher 1997). However, if there are emergent properties of multiple mutualists, the selective effects of mutualists will be nonadditive. In this case, selection exerted by the mutualist community on the focal species should not be predictable based simply on selection imposed by an individual partner species (Strauss & Irwin 2004). Imagine for example, that two separate mutualist species each provide a resource (e.g., N, P, H_2_0, defense) that improves plant fitness compared to not having them, but that fitness is highest when interacting with both mutualists. In this instance, the traits or trait combinations leading to highest fitness when interacting with either mutualist alone, both, or neither are likely to differ. Surprisingly few studies have explicitly measured selection generated by multispecies mutualisms (e.g., Sahli & Conner 2011), especially among mutualists conferring different types of rewards. Thus, it is difficult to assess the frequency and strength of nonadditive selection resulting from these common but complex interactions.

Multiple mutualists can also influence evolution in mutualisms by affecting fitness alignment, the correlation in fitness functions, between interacting mutualists (Jones *et al.* 2015). The interaction between species is designated a mutualism based on a comparison to when one partner is absent (Fig. 1A). However, given that an interaction is a mutualism, an important question is whether fitness interests are aligned between partners. Does higher fitness for one partner result in higher fitness for the other in a straight-forward way (Fig. 1B*(ii)*)? It is also unknown how additional mutualists alter the fitness alignment between a pair of interacting mutualists (Fig. 1B*(i, iii)*). For example, is the relationship between plant and N-fixing bacterial fitness altered by the presence or absence of mycorrhizal fungi? The fitness alignment between mutualist partners will also affect the rate and strength of positive feedbacks within the component interactions. In the hypothetical scenario shown in Fig. 1A and 1B*(ii)*, plant and rhizobial fitness are tightly correlated in the absence of mycorrhizal fungi, and this correlation is strengthened by the presence of mycorrhizal fungi (Fig. 1B*(i)*). All else being equal, we would expect combinations leading to highest fitness of both partners to have an advantage. In contrast, in another hypothetical scenario for the relationship between plant and rhizobial fitness in the presence of mycorrhizal fungi (Fig. 1B*(iii)*), there are many plant and rhizobial combinations that lead to nearly equivalent plant fitness yet varying bacterial fitness. The potential for additional mutualists to modify the fitness alignment between a pair of mutualist partners is an essentially unaddressed empirical question.

**Figure 1:**
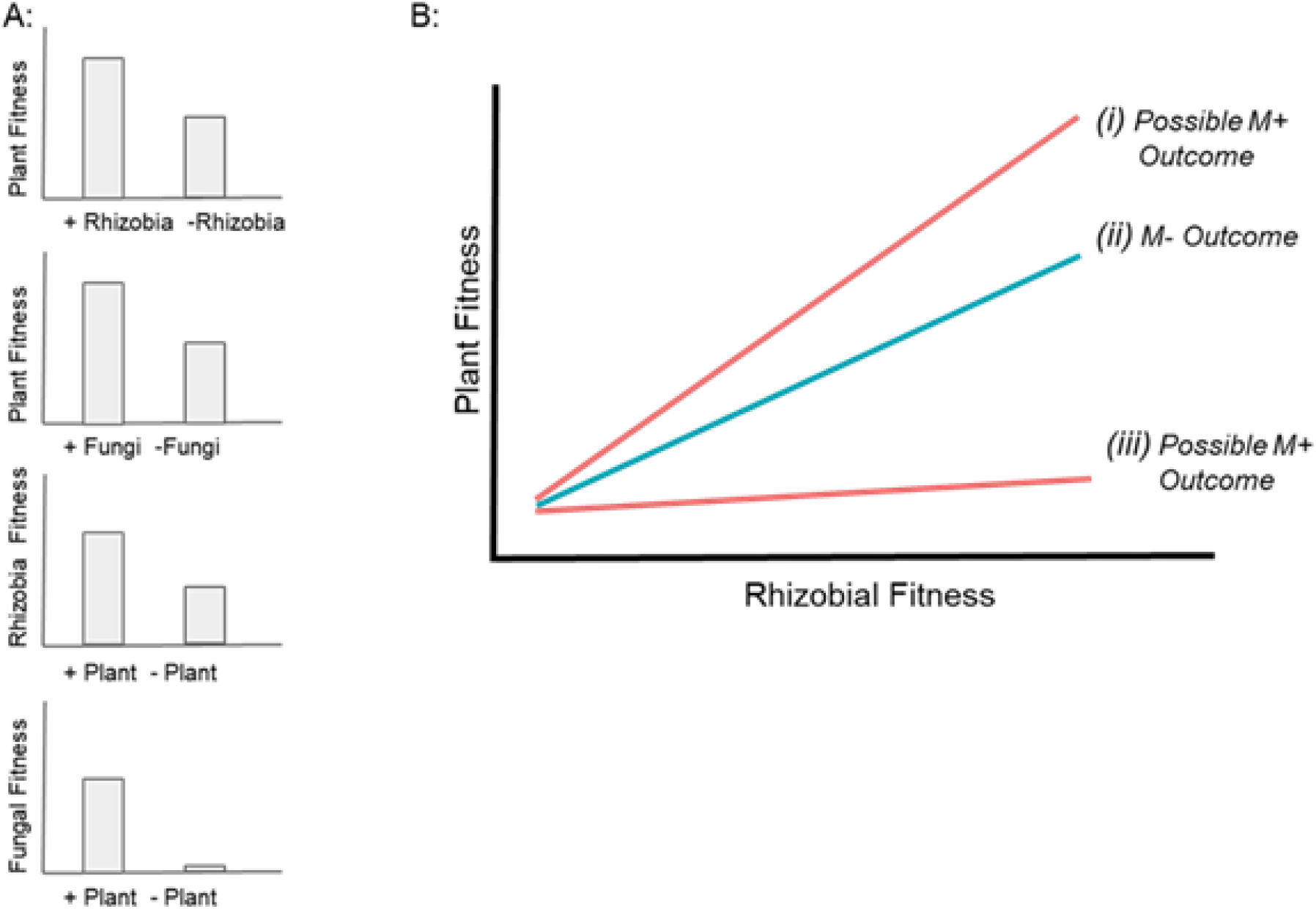
Schematic showing multiple mutualisms and fitness alignment. In (**A**), plants have higher fitness with rhizobia and fungi than without, and rhizobia and fungi have higher fitness with plants than without (i.e., the relationships are mutualisms). In (**B**) we show a two hypothetical case where the presence of mycorrhizal fungi alters the relationship between bacterial and plant fitness. *(ii)* shows the relationship in the absence of mycorrhizal fungi (M−), *(i)* shows case where mycorrhizal fungi strengthens fitness alignment between the plant and bacteria via complementarity of microbial mutualist rewards, and *(iii)* shows case where carbon demands of two mutualists lead to a weaker return for plants of rhizobial fitness.

To address the role of MMEs in the eco-evolutionary dynamics of mutualisms, we grew replicate plants from 213 genotypes of *Medicago truncatula* in a 2×2 factorial experiment manipulating the presence of rhizobia and mycorrhizal fungi. Specifically, we asked two related questions: (1) Do third-party mutualists (i.e. the multiple mutualist context) enhance or diminish fitness alignment? and (2) Do multiple mutualists alter heritability of and impose nonadditive selection on host traits?

## Methods

### Study System

We used the tripartite mutualism between the annual legume *Medicago truncatula* (‘Barrel Medic’), rhizobia (*Ensifer meliloti* Rm1021), and mycorrhizal fungi (*Rhizophagus irregularis* DAOM197198) to investigate multiple mutualists effects on selection and fitness alignment. We chose *M. truncatula* because: (i) it interacts with two common plant mutualists (rhizobia and mycorrhizal fungi) that primarily provide distinct rewards to plants (fixed N and soil P/water, respectively) and are tractable to manipulate (Afkhami & Stinchcombe 2016; Kafle *et al.* 2019), (ii) plant genotypes vary in their response to multiple mutualists (Franklin *et al.* 2020), and (iii) it has a hapmap project with >200 distinct plant genotypes available from across its range (http://www.medicagohapmap.org). These genotypes allow for inclusion of a wide range of host genetic diversity and thus more rigorous genotypic selection and fitness alignment analyses.

### Experiment

#### Experimental design

We grew 213 *M. truncatula* genotypes (Table S1) in four microbial environments: no microbes (M−R−), rhizobia alone (M−R+), mycorrhizal fungi alone (M+R−), and both microbes (R+M+) using a completely randomized block design with five spatial blocks. For each genotype, we mechanically scarified and surface-sterilized ~25 seeds in a bleach solution, rinsed with sterile water, and germinated them on sterile 0.8% water agar plates for ~36hrs at 4°C in the dark followed by ~18 hrs at 22°C (Garcia *et al.* 2006; Afkhami & Stinchcombe 2016). We planted germinants into 164ml cone-tainers (Stuewe and Sons, Oregon) filled with sand. Before planting, we sterilized all pots and sand at 121°C three times (45 minute cycle) and then inoculated with appropriate microbial treatments one and two weeks after planting to encourage colonization and nodulation. Bacterial inoculum was grown for 36hrs in TY media and diluted to ~10^6^ cells/ml (OD600 = 0.1) with ddH2O (Simonsen & Stinchcombe 2014). Germinants in R+ pots received 1 ml of inoculant and those in R- pots received the same ‘inoculant’ without rhizobia. Mycorrhizal inoculum for M+ germinants was ~300 spores in sterile water (Premier Tech, Riviere-du-Loup, Quebec, Canada; Antunes *et al.* 2008; Powell *et al.* 2009) while pots with M- germinants received inviable inoculum (achieved by autoclaving spores four times for 45 min at 121 °C). We grew plants in the greenhouse until fruit production (~8 months) with supplemental lighting to reflect daylength in *M. truncatula’s* native range and fertilizer supplied at 3 month intervals (1:1:1 ppm N:P:K). In total, we planted ~4260 seeds into pots (=4 microbial treatments x 213 genotypes X 5 replicates).

#### Data collection and harvest

We collected and counted mature *M. truncatula* pods (fruits) as they were produced. At harvest, we counted branches (Moreau 2006) and collected above-ground biomass, which we dried at 60°C, and then weighed. Data on belowground-biomass and rhizobia nodule number were collected for a subset of plants in each genotype-microbial environment combination (n=323.5±24.3 and n=314.5±10.5 per treatment, respectively; recall that we cannot count nodules in R- treatments). We calculated the ratio of root to shoot mass for each plant (to examine allocation) and genotype means for branches, root:shoot mass, pod number, and nodule number in each microbial environment separately. We assigned a fitness of zero to individuals who did not produce mature pods.

### Data Analyses

Analyses were conducted in R (version 3.6.1) using the car and boot packages for AN(C)OVA and bootstrapping (Fox & Weisberg 2018; Canty & Ripley 2019).

#### Test of main effects and genetic variation

To examine whether plant genetic variation and microbial context affected plant fitness, we first used an ANOVA with pod number as a response variable and explanatory variables of mycorrhizal fungi (presence/absence), rhizobia (presence/absence), plant genotype, and all interactions among these variables. We square-root+1 transformed pod number prior to analysis to improve normality. We conducted the same analysis for root:shoot ratio (square root+1 transformed) and branch number. These two traits were selected based on preliminary analysis that found other plant size traits were strongly correlated with branch number while root:shoot was uncorrelated with these traits (Table S2). For plants inoculated with rhizobia (M+R+ and M−R+ treatments), we also examined whether plant genetic variation and the presence of a third-party mutualist impacted nodule number (proxy for rhizobia fitness) using an ANOVA with explanatory variables of mycorrhizal fungi presence/absence, plant genotype, and their interaction. In all models, we included a block term to account for spatial variation and used type III sum of squares. We considered block and genotype to be fixed effects because they did not represent random samples of spatial or genetic variation, respectively, about which we wished to generalize (Newman *et al.* 1997).

As a complement to our tests of main effects of treatments and genotypes, we also estimated and compared the heritability of plant traits in each microbial environment. Briefly, and only for estimating heritability, we fit mixed-model ANOVAs for traits with block as a fixed effect and line as a random effect for each microbial environment separately, using restricted maximum likelihood. We estimated broad sense heritability, H^2^, as *V*_*line*_/(*V*_*line*_ + *V*_*residual*_). To characterize uncertainty in heritabilities, we adapted a procedure described by (Houle & Meyer 2015) for G matrices. We considered REML estimates of *V*_*line*_ and *V*_*residual*_ as means of a bivariate normal distribution, with a covariance matrix equal to the asymptotic variance matrix of the REML estimates. We drew 10,000 values from these bivariate normal distributions, and estimated confidence limits from the 2.5th and 97.5th percentiles of these values. We estimated heritability and its corresponding uncertainty in SAS v9.4, using Proc Mixed and IML.

#### Do third party mutualists enhance or diminish fitness alignment?

To assess whether mycorrhizal fungi affected fitness alignment between rhizobia and host plants, we used an ANCOVA with a response variable of mean pod number (square root + 1 transformed) and a fixed effect of mycorrhizal fungi (presence/absence), a continuous predictor variable of nodule number, and their interaction. A significant interaction between fungi presence and rhizobia fitness indicates an effect of a third-party mutualist on fitness alignment of the rhizobia-plant mutualism. We followed up on a significant interaction with separate regressions for M+R+ and M−R+ plants to assess fitness relationships in each microbial environment.

#### Do multiple mutualists impose nonadditive selection on host traits?

To determine how multiple mutualists select on their shared host, we used genotypic selection analysis (Rausher 1992). We used ANCOVA to determine how each microbe individually and jointly altered the relationship between plant traits and fitness (i.e., altered the strength and/or direction of selection gradients). We modeled relative plant fitness using fixed factors of rhizobia (R+/R−) and mycorrhizal fungi (M+/M−), two traits (branching and root-to-shoot mass ratio), the interaction between microbes, and the interactions between microbes and traits. We calculated plant relative fitness for this analysis by dividing mean fitness for each genotype by the overall mean fitness across all environments. While doing so implicitly invokes a hard selection model (De Lisle and Svennson 2017), we made this choice so that differences in mean fitness and trait values, which were of *a priori* interest, were preserved in our ANCOVA models (see Batstone et al. 2020). To verify that any significant interaction terms were not driven solely by differences in mean fitness between treatments, we repeated this analysis with plant relative fitness calculated within a treatment, and found similar statistical results for this ANCOVA (Table S7), including the same significance for the three-way interactions. A significant three-way interaction between rhizobia, fungi, and a trait in this analysis indicates nonadditive selection on that trait. We followed up on significant three-way interactions with multiple linear regression of relative plant fitness (calculated across treatments) on the traits in each microbial environment separately to determine the selection gradient in each environment. Bias corrected 95% CIs around each selection gradient were calculated using 10,000 bootstraps.

## Results

### Overview of main effects and genetic variation

Pod number (a measure of host plant absolute fitness) varied significantly among genotypes (genotype main effect: F_212,3397_=4.55, P<<0.00001; Table S3), indicating genetic variation for plant fitness. Microbial context also altered host fitness, but the strength, direction, and the nonaddivitity of these effects depended on host genotype (e.g., rhizobia×fungi×genotype interaction: F_212,3397_=1.163, P=0.059; Table S3). Nodule number (a measure of rhizobial absolute fitness) varied significantly by plant genotype (F_155,251_=2.84, P<<0.00001; Table S3). Mycorrhizal fungi also affected nodule production with genotypes varying in their response to the third-party mutualists, indicating genetic variation among hosts mediate mycorrhizal fungi’s effect on nodulation (fungi×genotype interaction: F_155,251_=1.33, P=0.024; Table S3). We found that host genotype (F_138,880_=5.10, P<<0.00001; Table S3) and the interaction between mycorrhizal fungi, rhizobia, and host genotype (F_138,880_=1.229, P=0.0482) significantly affected the branch number trait, and that investment in roots versus shoots (root:shoot ratio) differed among plant genotypes (F_112,398_=1.72, P<<0.00001; Table S3) and depended on microbial environment (rhizobia×fungi: F_1,398_=3.57, P=0.0595).

Microbial treatments affected broad sense heritability of plant traits, but not in a consistent way across traits (Table S4). For example, broad sense heritability of pod number, was highest in the treatment with both mutualists (H^2^ = 0.35), and lower in the three remaining treatments (H^2^ ranges 0.12 to 0.16). For branch number, heritability was highest with both mutualists (H^2^ = 0.51), followed by rhizobia only (H^2^ = 0.42), followed by the treatments lacking rhizobia (H^2^ = 0.18 and 0.26 for with and without fungi, respectively). In contrast for nodules (which require the presence of rhizobia), heritability was higher in the absence of mycorrhizal fungi (H^2^ = 0.48) than in their presence (H^2^ = 0.29), though the uncertainty estimates of these heritabilities overlap.

### Do third party mutualists enhance or diminish fitness alignment?

Not only did third-party mutualists mycorrhizal fungi impact the relationship between rhizobia and the plant host, but surprisingly the fungus had a strong impact on fitness alignment (fungi presence × rhizobia fitness: F_1,355_=4.94, P=0.0269; Fig. 2, Table S5a). In absence of mycorrhizal fungi, there was a weak, but significant, positive relationship between rhizobia relative fitness (calculated from nodule number) and plant relative fitness (calculated from fruit number) (F_1,173_=6.9, P=0.0094; Fig. 2, Supplementary Table S5b). In the presence of mycorrhizal fungi, rhizobia and plant fitness were substantially more positively aligned (F_1,182_=21.45, P<0.00001; Fig. 2, Table S5b) with a slope that is 2.5 times greater than when the third-party mutualist was absent.

**Figure 2.**
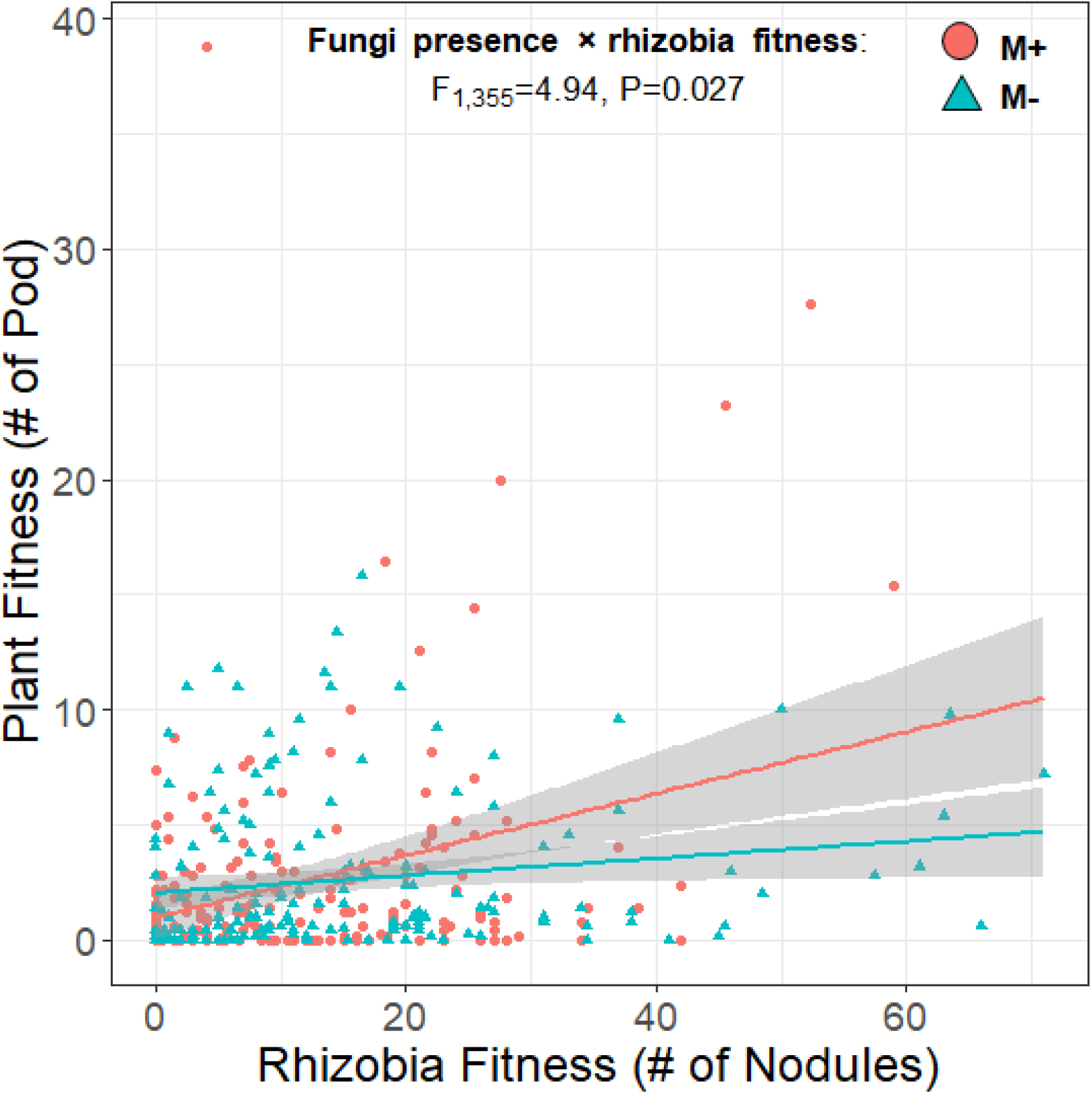
Fitness alignment between the rhizobia and the host plant was stronger in the presence of mycorrhizal fungi. Each point represents an individual plant genotype, and lines indicate relationship between plant and rhizobia fitness in each mycorrhizal treatment environment. Y-axis value is the mean fitness of a plant genotype as indicated by fruit production (pod number), and x-axis value is the mean fitness of the rhizobia associating with that plant genotype as indicated by nodule number. The red circles and line represent the treatment with mycorrhizal fungi and blue triangles and line represent the treatment without mycorrhizal fungi.

### Do multiple mutualists impose nonadditive selection on host traits?

Rhizobia and mycorrhizal fungi exerted significant nonadditive selection on branching of their host plant (fungi × rhizobia × branch number: F_1,664_= 5.71, P=0.0171; Fig. 3, Table S6-7). In the absence of all microbes or in the presence of just mycorrhizal fungi, selection on plant branch number was very weak (β_M−R−_=0.031 and β_M+R−_=0.034; Fig. 3, Tables 1, S8). In the presence of rhizobia (alone), we observed a positive relationship between branch number and relative plant fitness, indicating selection for increased branching of host plants occurs in the presence of rhizobia (β_M−R+_=0.278, Tables 1, S8). Interestingly, the strength of selection on branching doubled in the presence of multiple mutualists (β_M+R+_=0.557) compared to plants grown with rhizobia alone and was >16x stronger than with mycorrhizal fungi alone. The selection gradient in the presence of both partners (0.557) is nearly double the expected additive selection gradient of 0.281 ; calculated following TerHorst et al. (2015)(β_additive_=0.034+0.278-0.031). The nonadditive selection gradient, which quantifies how much selection is modified by indirect ecological interactions, was 0.276 (calculated as β_M+R+_ - β_additive_). Selection gradients reported here are from the multiple linear regression (Table 1), but univariate analyses (where each trait’s relationship with plant fitness is analyzed separately) showed nearly identical values (Table S8).

**Figure 3.**
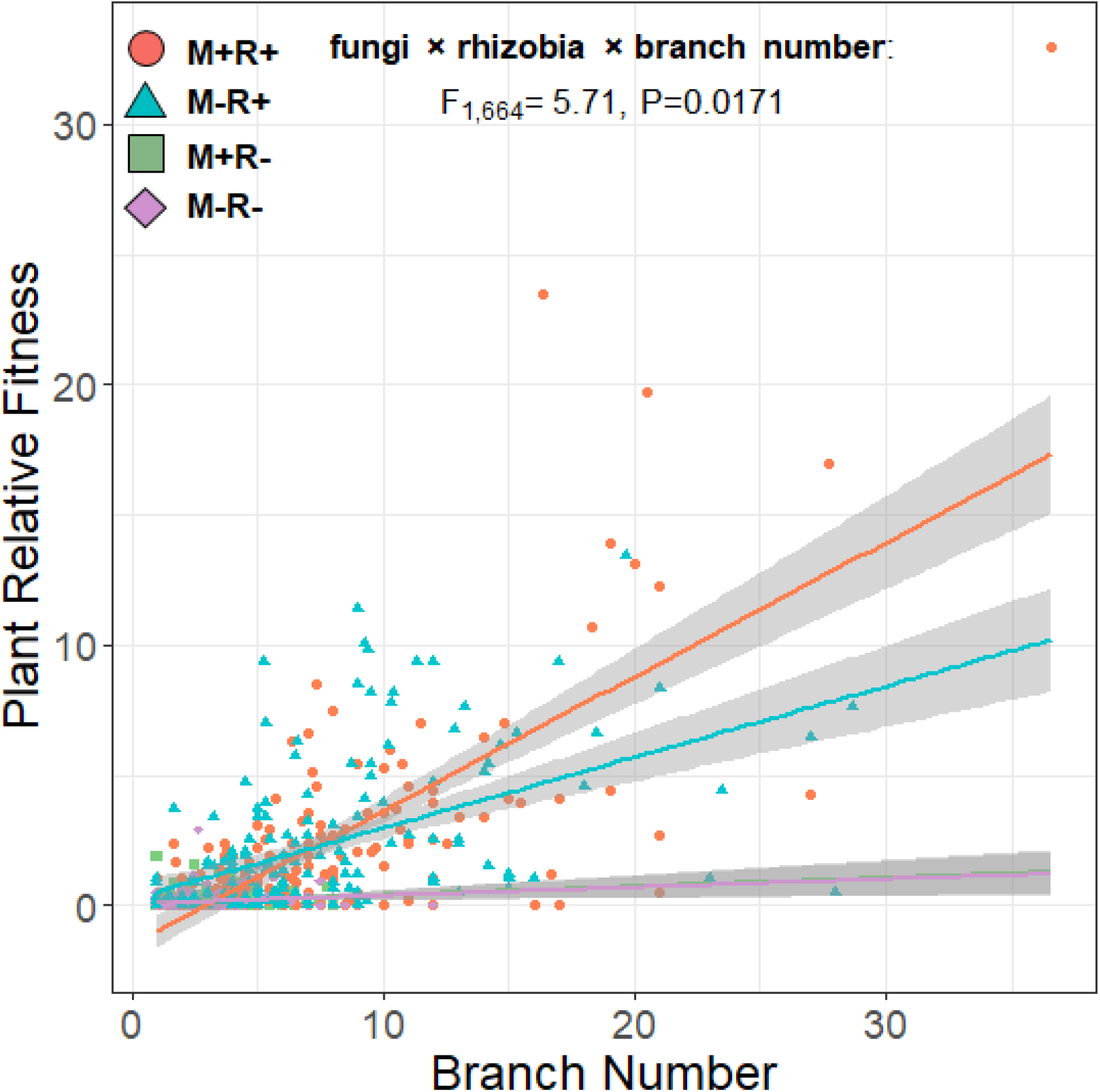
Mycorrhizal fungi and rhizobia select nonadditively on their host plant’s branching. Each point represents an individual plant genotype, and lines indicate relationships between mean branch number and plant relative fitness in the four microbial treatment environments. The purple diamonds and line represent the no microbe treatment, green squares are the mycorrhizal fungi only treatment, blue triangles are the rhizobia only treatment, and red circles are the multiple mutualist (both microbe) treatment. Note that the green line is not visible because the slopes of M−R− and M+R− lines (purple and green) are nearly identical.

**Table 1.**
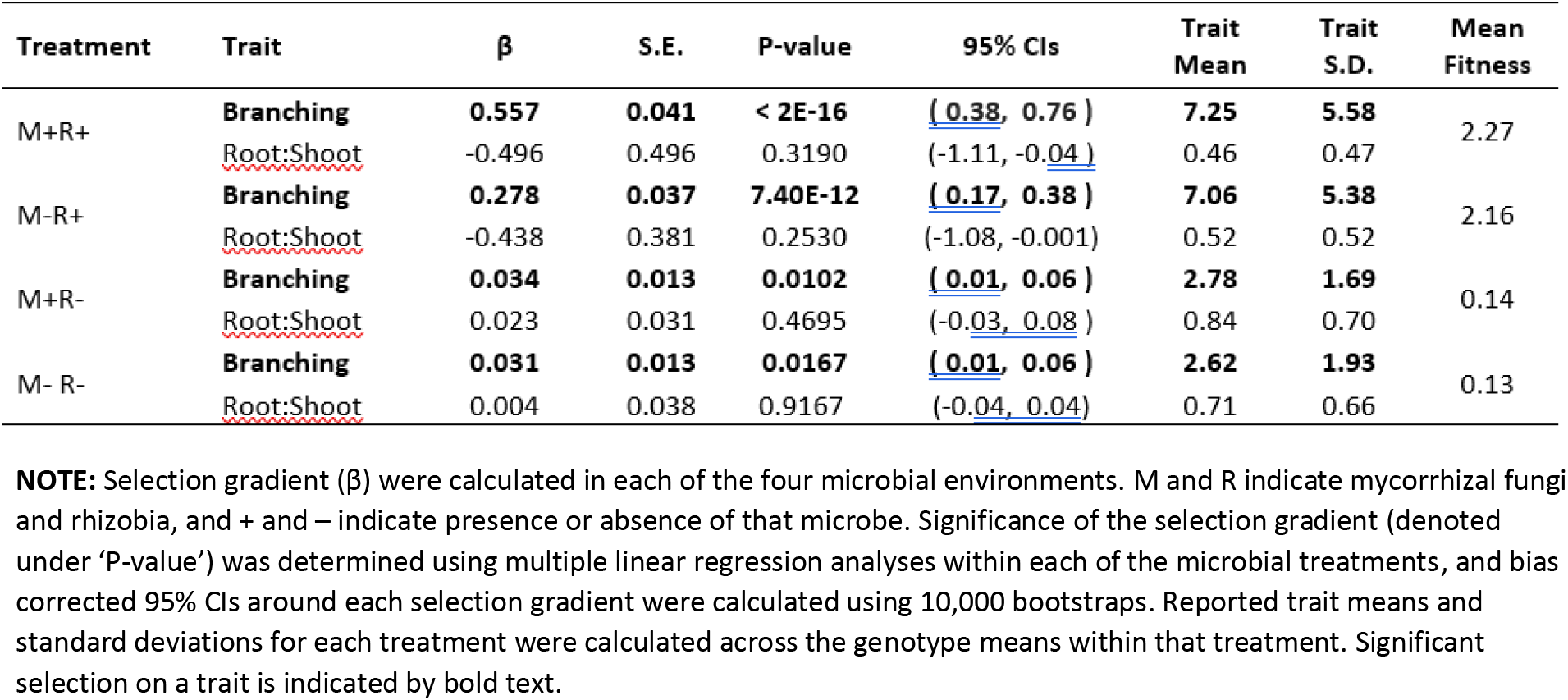
Results of analyses to determine selection gradients for microbial effects on plant traits.

## Discussion

A large gap remains in our understanding of how multispecies mutualistic interactions impact evolutionary trajectories of species and their interactions. Our study revealed two key findings: We showed for the first time that (1) multispecies mutualisms not only can have non-additive effects on performance, but also on selection and heritability of host traits and (2) third-party mutualists can enhance fitness alignment between mutualist species. Below, we first discuss the implications of non-additivity of fitness, heritability, and selection in multispecies mutualisms. We then discuss synergism and conflict in the direction of selection in multispecies mutualisms and fitness alignment between mutualists.

### Non-additivity of performance, fitness, heritability and selection

Multiple mutualist interactions can have non-additive effects on performance traits and fitness components that could affect the demographics and evolutionary trajectories of legumes. In past studies of *Medicago truncatula*, we showed that there were non-additive effects on overall plant performance (biomass), on gene expression profiles, and on coexpression network structure (Afkhami & Stinchcombe 2016; Palakurty *et al.* 2018; Hernandez *et al.* 2020). Here, we detected non-additive effects of multiple mutualists on plant fitness and an architectural trait, branch number. These non-additive effects thus have the potential to affect plant population dynamics, through fecundity, and the interaction between plants (as branchier plants may shade each other more). Taken together, the pervasive multiple mutualist effects that we have demonstrated suggest that understanding plant traits--including gene expression, size, shape, fecundity-- will require embracing the effects of multiple mutualists simultaneously, rather than in pairs. Whether MMEs have evolutionary consequences, however, had been relatively unexplored.

Our data convincingly show that multiple mutualist effects and non-additivity extend to the key components of microevolutionary change, genetic variation and natural selection. Pod number, which is a fitness component in annual plants with high selfing, showed the highest heritability in the presence of multiple mutualists, with a value two to three-fold higher than with single mutualists (H^2^ = 0.35, versus 0.12 and 0.16 for mycorrhizal and rhizobia-only treatments). While differences in heritability can be driven by changes in environmental variance, we also observed substantial differences in genetic variance components (Vg) and coefficients of genetic variation (Table S4), indicating that increased evolutionary potential in the presence of multiple mutualists was neither driven by reduced environmental variance nor solely by increases in mean pod number. On a practical level, higher heritabilities suggest that artificial selection and planned breeding programs for legumes (of which there are many important food crops; Graham & Vance 2003) would expose more genetic variation and lead to larger responses to selection in the presence of multiple mutualists. On a conceptual level, these data also suggest substantially more genetic variation for fitness components in the presence of multiple mutualists. While acknowledging that pod number may not represent *true fitness* and that our study was carried out in the greenhouse, these results nonetheless suggest that MMEs could affect the rate of adaptation, which will be determined by genetic variation for fitness (Fisher 1930). Empirically testing this in the MME framework in the field would be challenging, though recent work (Kulbaba *et al.* 2019; Sheth *et al.* 2018) suggests some ways forward.

Beyond potential effects on the overall rate of adaptation, genetic variance in and selection on branch number was affected by MMEs, which could impact evolution of this trait. Heritability for branch number was significantly elevated when both mutualists were present (as were genetic variances and coefficients of genetic variation), and branch number was subject to significant non-additive selection in the presence of both mutualists. Because of increased heritability and genetic variance in branch number in the presence of both mutualists coupled with stronger selection, any predicted evolutionary response of this trait is much greater than in conditions with one or no mutualists. The evolutionary response of a single trait can be predicted from the univariate version of Lande’s equation, R = Vgβ, where R is the response to selection, Vg is the genetic variance, and β represents the selection gradient for a trait. Solving this equation for branch number suggests negligible changes in branch number in the absence of rhizobia (0.03 to 0.04 branches), moderate changes in the M−R+ treatment (~4.4 branches), and nearly a 3-fold larger evolutionary response in the M+R+ treatment (~12.25 branches).

While these are almost certainly over-estimates (mean branch number in these treatments ranged from 2.6 ± 0.11 to 7.3 ± 0.33; 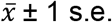), they nonetheless illustrate that MMEs have substantial quantitative effects on predicted microevolutionary responses. Much effort has focused on the ecological conditions that can produce non-additive selection (Haloin & Strauss 2008; TerHorst *et al.* 2015), and how ecological interactions in complex communities can alter the strength of selection in ways that cannot be predicted from pairwise experiments. Our results suggest that this holds for the expression of genetic variation, and that combined with non-additive selection, could dramatically affect microevolutionary trajectories.

One potential caveat to our investigation of how MMEs affect heritability and genetic variances is that we performed univariate analyses, which preclude estimates of genetic covariances between traits. (We note that our estimates of selection are from multiple regression, and thus account for indirect selection on other included traits). A multivariate genetic approach would provide information about genetic covariances between traits as in (Ossler & Heath 2018)-- for example, between branch number and root:shoot ratio, and whether multiple mutualists affected **G** matrices. We elected not to pursue this approach because of difficulties in making comparisons among **G** matrices separately estimated in four separate experimental treatments. Given evidence of MMEs affecting plant performance, phenotypes, molecular traits, genetic variances, and selection, it seems likely that they can also affect **G**, although this remains untested.

### Selection in Multiple Mutualisms

Much of the work on selection in multispecies interactions has focused on conflicting selection, especially between antagonistic and mutualistic interactions (Gómez 2003; Kessler & Halitschke 2009; Pérez-Barrales *et al.* 2013) and among multiple antagonistic interactions (e.g., Miller *et al.* 2014). However, concordant selection (i.e. agreement in the direction of selection) has also been observed. For instance, Sletvold *et al.* (2015) found that the same pollinator and herbivore that imposed conflicting selection on one trait (flowering time), imposed concordant selection on another trait (nectar spurs). The few previous multispecies mutualism selection studies have also typically been motivated by identifying conflicting selection generated by different partners, as this could either constrain evolution in the shared partner or lead to adaptive differentiation, depending on the trait’s genetic architecture (Assis *et al.* 2020). These studies also predominantly focused on within-guild multiple mutualisms (i.e. partner species provide the same resource to the host), where partner mutualists may compete to interact with a shared host. In particular, studies of plant-pollinator interactions have often documented conflicting selection. For instance, Kulbaba & Worley (2013) found conflicting selection on corolla diameter by hummingbirds and hawkmoths. Similarly, floral corolla flare was predicted to be under selection to be narrower with hummingbirds but predicted selection was neutral with bumblebees (Aigner 2004). However, multiple pollinator mutualists have been shown to impose concordant selection on some traits; Sahli & Conner (2011; re-analysis by TerHost et al. 2015), for example, documented concordant selection on the degree of anther exertion by sweat bees and larger bees. Since floral phenotypes typically determine the ability of mutualists to interact via matching, these traits may be predicted to more often be under conflicting selection than traits involved in resource-exchange mutualisms, where concordant selection -- and even synergistic concordant selection -- may play a more important role.

We know of no other cases of synergistic (concordant) selection in a multispecies mutualism, begging the questions of when it should be expected and why it has not been documented previously. Synergistic selection may not have been previously shown because most studies of multispecies mutualisms simply do not measure selection (Mack & Rudgers 2008; Palmer *et al.* 2010; Orivel *et al.* 2017) and those that do are usually measure selection in a pairwise fashion (i.e, selection imposed by each species individually, but not collectively, or as a group but not individually) and thus are not designed to test nonadditivity (TerHorst *et al.* 2015). A notable exception is Sahli & Conner (2011), who estimated how pollinators individually and as a group imposed selection on floral morphology. However, unlike our study which demonstrated synergistic selection on branch number, re-analysis of their data by TerHorst et al. (2015) found evidence for concordant, but not synergistic, selection.

We posit two factors may lead to synergistic selection: the importance of the trait in acquiring resources later traded to mutualist partners (e.g., traits involved in carbon acquisition, when C-based rewards are provided to microbial partners) and complementarity of partner-provided rewards (Afkhami *et al.* 2014, 2020). For host resource-acquisition traits, we expect that they will be subject to synergistic selection when the same resource is provided to all partners: in these cases, there should be selection favoring trait values that increase the size of the resource ‘pie.’ In these cases, while selection may act in the same direction on traits when only one mutualist is present, environments with multiple mutualists may lead to an additive increase in selection on those traits. The nonadditivity of selection in our study likely resulted from complementarity of the rewards provided to host plants by microbial partners. In the absence of rhizobia, plants in our experiment did not perform well and produced very few pods (a good estimate of fitness for the *M. truncatula;* Friesen *et al.* 2014) and relatively few branches, regardless of whether mycorrhizal fungi was present. Without fixed N from rhizobia, plant growth and reproduction was limited, such that increased branching could not improve plant fitness through increased carbon acquisition or allocation to mycorrhizal fungi. In the presence of both partners, selection was stronger than what was observed when rhizobia was present alone, likely because rhizobia and mycorrhizal fungi alleviated different limitations on growth and reproduction. While complementarity of partner-conferred rewards increases the probability of synergistic selection, even functionally redundant mutualists that provide the exact same benefit may result in synergistic selection on traits if each partner alone would not provide sufficient resources to allow host survival and reproduction. Alternatively, in the case of saturating effects of the benefit received on the trait (or fitness), we would predict that multiple mutualists would result in non-additive selection, where both partners still produce selection in the same direction (i.e., there is no conflicting selection) but the total strength of selection with multiple mutualists is lower than additive predictions.

### Fitness Alignment in Mutualisms

A classic expectation in mutualisms is that they should be subject to “cheating” that could threaten their persistence and stability (Ferriere *et al.* 2002; Douglas 2008). The widespread expectation of cheating has led to a large literature on mechanisms that can maintain mutualisms in the face of cheating (e.g., partner choice, partner fidelity feedback, screening, and sanctions), definitions of cheating (e.g., Ghoul *et al.* 2014; Jones *et al.* 2015), and whether there is any evidence of cheating (Frederickson 2013). One important mechanism capable of stabilizing mutualisms is partner fidelity feedback, where mutualisms persist because of positive feedbacks between partners: symbionts benefit by improving host fitness, and hosts benefit by improving symbiont fitness (Sachs *et al.* 2004; Weyl et al. 2010). A key prediction of this view is a positive correlation between host and symbiont fitnesses, exactly as we observed (Figure 2). Our data show that plant genotypes with high fitness also produce lots of nodules, a fitness component of their rhizobial partners.

Interestingly, while plant and rhizobia fitness are positively and significantly correlated when mycorrhizal fungi are absent, the correlation is appreciably stronger in their presence. In other words, a third-party mutualist leads to scenario Figure 1(i), by enhancing the fitness alignment of interacting species. As a consequence, if this is a general finding, it may be that multispecies mutualisms can increase the persistence of the component interactions by more tightly aligning the fitness of the interactors.

We suggest that synergistic effects of multiple mutualists are likely to facilitate partner fidelity feedbacks. The logic is straightforward: if having multiple mutualists improves host fitness beyond what would be expected from interacting with single mutualists, hosts would be in better than expected condition to provide fitness benefits to symbionts. While positive feedbacks are also expected in the additive case, the synergistic effects of multiple mutualisms would lead to the strongest feedbacks, while only conflicting selection would reduce them. Evaluating the generality of these results requires more fully characterizing the fitness alignment of multiple interacting species.

### Conclusions and Prospects

Multiple mutualists can have significant, synergistic effects on heritability and selection for host traits, and can substantially influence fitness alignment within mutualisms. Taken together with the well-established fact that organisms often interact with multiple mutualists (Afkhami *et al.* 2014, 2020), these outcomes demonstrate the importance of multiple mutualist effects for evolutionary trajectories of organisms and their beneficial interactions. In our opinion, there are at least three types of studies that will be valuable for furthering our understanding of multispecies mutualisms’ roles in eco-evolutionary dynamics. First, in addition to considering host traits, future work should include the effects of mutualists on traits in all partners (e.g., do rhizobia affect mycorrhizal fungi’s allocation to intra- vs extraradical hyphae; van Aarle & Olsson 2008), rather than remaining host-centric. Second, more studies testing how multispecies beneficial interactions impact selection and heritability of host traits across a wide range of systems is crucial to determine the generality of synergistic and conflicting selection. Field common garden studies with tractable partners would be especially valuable. Third, to link complementarity of partner mutualists (Afkhami *et al.* 2014, 2020) with synergistic selection, studies factorially manipulating within- versus across-guild multiple mutualisms and measuring selection and complementarity are needed. Collectively, studies like these will make meaningful progress in understanding the eco-evolutionary consequences of multispecies mutualisms.

## Acknowledgements

We thank N. Aryan for help with data collection and the Medicago Hapmap Project for providing seeds. Our work was supported by University of Toronto EEB Postdoctoral Fellowship, NSF IOS-1401840, NSF DEB-1922521, and NSF DEB-2030060 to MEA and an NSERC Discovery Grant to JRS. MLF acknowledges support from NSF DEB-1823419, NSF DEB-1943628, and the USDA National Institute of Food and Agriculture, Hatch project 1014527.

## Notes

### Competing Interest Statement

The authors have declared no competing interest.

