## Supplementary Materials for "Multiple Mutualist Effects generate synergistic selection and strengthen fitness alignment in a tripartite interaction between legumes, rhizobia, and mycorrhizal fungi"

**<u>Supplementary Table S1</u>: Genotypes included in this study.**

| | | | | | | | |
|---|---|---|---|---|---|---|---|
| HM001 | HM043 | HM073 | HM109 | HM149 | HM182 | HM259 | HM314 |
| HM002 | HM044 | HM074 | HM111 | HM150 | HM183 | HM260 | HM315 |
| HM003 | HM045 | HM075 | HM112 | HM151 | HM184 | HM266 | HM316 |
| HM004 | HM046 | HM076 | HM114 | HM152 | HM185 | HM267 | |
| HM005 | HM047 | HM077 | HM115 | HM153 | HM186 | HM268 | |
| HM006 | HM048 | HM078 | HM117 | HM154 | HM187 | HM269 | |
| HM007 | HM049 | HM079 | HM118 | HM155 | HM188 | HM270 | |
| HM008 | HM050 | HM080 | HM119 | HM156 | HM189 | HM271 | |
| HM009 | HM051 | HM081 | HM120 | HM157 | HM190 | HM276 | |
| HM010 | HM052 | HM082 | HM121 | HM159 | HM191 | HM277 | |
| HM011 | HM053 | HM083 | HM122 | HM160 | HM192 | HM279 | |
| HM013 | HM054 | HM084 | HM124 | HM161 | HM193 | HM287 | |
| HM014 | HM055 | HM085 | HM125 | HM162 | HM194 | HM288 | |
| HM015 | HM056 | HM086 | HM126 | HM163 | HM195 | HM289 | |
| HM016 | HM057 | HM087 | HM127 | HM164 | HM196 | HM290 | |
| HM026 | HM058 | HM088 | HM128 | HM165 | HM197 | HM293 | |
| HM027 | HM059 | HM089 | HM129 | HM166 | HM198 | HM294 | |
| HM028 | HM060 | HM091 | HM130 | HM167 | HM199 | HM296 | |
| HM031 | HM061 | HM092 | HM131 | HM168 | HM200 | HM297 | |
| HM032 | HM062 | HM093 | HM133 | HM169 | HM201 | HM298 | |
| HM033 | HM063 | HM095 | HM134 | HM170 | HM202 | HM299 | |
| HM034 | HM064 | HM096 | HM135 | HM172 | HM203 | HM301 | |
| HM035 | HM065 | HM097 | HM138 | HM173 | HM205 | HM304 | |
| HM036 | HM066 | HM098 | HM139 | HM175 | HM206 | HM305 | |
| HM037 | HM067 | HM099 | HM141 | HM176 | HM207 | HM307 | |
| HM038 | HM068 | HM101 | HM143 | HM177 | HM208 | HM308 | |
| HM039 | HM069 | HM105 | HM145 | HM178 | HM209 | HM309 | |
| HM040 | HM070 | HM106 | HM146 | HM179 | HM251 | HM310 | |
| HM041 | HM071 | HM107 | HM147 | HM180 | HM253 | HM311 | |
| HM042 | HM072 | HM108 | HM148 | HM181 | HM256 | HM312 | |

**Supplementary Table S2: Correlations between measured plant traits.**

| | Branch | Aboveground Mass | Belowground Mass | Root:Shoot | Pod | Leaves at 6m | Leaves at 3m | Nodules |
|---|---|---|---|---|---|---|---|---|
| Branch | 1 | 0.88 | 0.63 | -0.09 | 0.66 | 0.81 | 0.38 | 0.44 |
| Aboveground Mass | 0.88 | 1 | 0.62 | -0.17 | 0.62 | 0.76 | 0.28 | 0.38 |
| Belowground Mass | 0.63 | 0.62 | 1 | 0.23 | 0.42 | 0.52 | 0.19 | 0.53 |
| Root:Shoot | -0.09 | -0.17 | 0.23 | 1 | -0.08 | -0.13 | -0.02 | 0.1 |
| Pod | 0.66 | 0.62 | 0.42 | -0.08 | 1 | 0.72 | 0.27 | 0.29 |
| Leaves at 6m | 0.81 | 0.76 | 0.52 | -0.13 | 0.72 | 1 | 0.34 | 0.41 |
| Leaves at 3m | 0.38 | 0.28 | 0.19 | -0.02 | 0.27 | 0.34 | 1 | 0.13 |
| Nodules | 0.44 | 0.38 | 0.53 | 0.1 | 0.29 | 0.41 | 0.13 | 1 |

**NOTE**: Correlations were calculated between all pairs of plant traits using their genotype means. 'Root:Shoot' was calculated as the ratio of above- to belowground biomass, 'Leaves at 6m and 3m' represent the number of leaves plant had after 6 months and 3 months of growth, respectively, and 'Nodules' represents the number of nodules at harvest. Analysis was done using the cor() function in R (version 3.6.1).

**Supplementary Table S3: Models testing effects of genotype, mycorrhizal fungi, rhizobia, and their interaction on plant and rhizobia fitness and plant traits.**

| Factor | Pod Number | | | Nodule Number | | | Branch Number | | | Root:Shoot | | |
|---|---|---|---|---|---|---|---|---|---|---|---|---|
| | DF | F | P | DF | F | P | DF | F | P | DF | F | P |
| Block | 4 | 4.900 | **0.0006** | 4 | 5.837 | **0.0002** | 4 | 1.176 | 0.3200 | 4 | 5.532 | **0.0002** |
| mycorrhizal fungi (myco) | 1 | 0.001 | 0.9728 | 1 | 4.720 | **0.0307** | 1 | 1.660 | 0.1979 | 1 | 0.852 | 0.3566 |
| rhizobia (rhizo) | 1 | 465.027 | **< 2.2E-16** | -- | -- | -- | 1 | 371.815 | **< 2E-16** | 1 | 25.260 | **7.6E-7** |
| Genotype | 212 | 4.546 | **< 2.2E-16** | 155 | 2.842 | **8.50E-14** | 138 | 5.098 | **< 2E-16** | 112 | 1.720 | **7.8E-5** |
| myco × rhizo | 1 | 0.047 | 0.8285 | -- | -- | -- | 1 | 0.720 | 0.3965 | 1 | 3.571 | **0.0595** |
| myco × genotype | 212 | 1.288 | **0.0040** | 155 | 1.326 | **0.0239** | 138 | 1.263 | **0.0299** | 112 | 0.885 | 0.7801 |
| rhizo × genotype | 212 | 3.497 | **< 2.2E-16** | -- | -- | -- | 138 | 3.131 | **< 2E-16** | 112 | 0.9757 | 0.5533 |
| myco × rhizo × genotype | 212 | 1.163 | **0.0585** | -- | -- | -- | 138 | 1.229 | **0.0482** | 112 | 0.6786 | 0.9926 |
| Residuals | 3397 | | | 251 | | | 880 | | | 398 | | |

**NOTE**: Treatment manipulating presence/absence of mycorrhizal fungi is indicated by "mycorrhizal fungi" ("myco"), treatment manipulating presence/absence of rhizobia is indicated by "rhizobia" ("rhizo"), and genotype of the plant is indicated as "genotype". Significant (and marginally significant) p-values are in bold. Note that analyses were conducted using ANOVA with a response variable of pod number and explanatory variables of mycorrhizal fungi (presence/absence), rhizobia (presence/absence), plant genotype, and all interactions among these variables, considering only genotypes for which there was data in every microbial treatment combination. Pod number and root:shoot were sqrt +1 transformed prior to analysis to improve normality, and analysis results were confirmed with permutational versions of these models using the aovp function in the lmPerm package.

**Supplementary Table S4: Genetic variation and heritability estimates for traits.**

| | Treatment | $V_g$ | $V_e$ | $H^2$ | 95% CIs | $\bar{X}$ | $\bar{X}$ SE | CVG |
|---|---|---|---|---|---|---|---|---|
| Pod | M+R+ | 14.6708 | 27.3491 | 0.3491 | (0.2780, 0.4125) | 2.2455 | 0.1977 | 1.7057 |
| | M+R- | 0.0455 | 0.3202 | 0.1244 | (0.0679,0.1794) | 0.1457 | 0.0185 | 1.4641 |
| | M-R+ | 4.9851 | 25.3675 | 0.1642 | (0.1044, 0.2212) | 2.2028 | 0.1701 | 1.0136 |
| | M-R- | 0.0528 | 0.3920 | 0.1186 | (0.0642, 0.1719) | 0.1365 | 0.0205 | 1.6822 |
| Trans Pod | M+R+ | 0.3273 | 0.7717 | 0.2978 | (0.2280, 0.3610) | 1.4677 | 0.0320 | 0.3898 |
| | M+R- | 0.0058 | 0.0338 | 0.1460 | (0.0875, 0.2022) | 1.0518 | 0.0061 | 0.0722 |
| | M-R+ | 0.1911 | 0.8003 | 0.1928 | (0.1305, 0.2513) | 1.4839 | 0.0307 | 0.2946 |
| | M-R- | 0.0050 | 0.0349 | 0.1262 | (0.0708, 0.1798) | 1.0471 | 0.0061 | 0.0678 |
| Branch | M+R+ | 22.8416 | 22.0254 | 0.5091 | (0.4127, 0.5911) | 7.3265 | 0.3340 | 0.6523 |
| | M+R- | 0.9625 | 4.4944 | 0.1764 | (0.0713, 0.2767) | 2.8409 | 0.1126 | 0.3453 |
| | M-R+ | 15.8877 | 22.1370 | 0.4178 | (0.2946, 0.5243) | 7.0965 | 0.3071 | 0.5617 |
| | M-R- | 1.3355 | 3.7652 | 0.2618 | (0.1360, 0.3758) | 2.6299 | 0.1083 | 0.4394 |
| Root:Shoot | M+R+ | 0.0225 | 0.4382 | 0.0489 | (0, 0.2009) | 0.5017 | 0.0436 | 0.2991 |
| | M+R- | 0.0882 | 1.0158 | 0.0799 | (0, 0.1861) | 0.8616 | 0.0565 | 0.3446 |
| | M-R+ | 0.0264 | 0.2839 | 0.0852 | (0, 0.3199) | 0.5050 | 0.0361 | 0.3219 |
| | M-R- | 0.0446 | 0.6078 | 0.0683 | (0, 0.2324) | 0.7015 | 0.0461 | 0.3009 |
| Trans Root:Shoot | M+R+ | 0.0038 | 0.0429 | 0.0820 | (0, 0.2348) | 1.2053 | 0.0139 | 0.0513 |
| | M+R- | 0.0123 | 0.0716 | 0.1461 | (0.0264, 0.2643) | 1.3312 | 0.0157 | 0.0831 |
| | M-R+ | 0.0046 | 0.0310 | 0.1282 | (0, 0.3400) | 1.2120 | 0.0122 | 0.0558 |
| | M-R- | 0.0058 | 0.0543 | 0.0970 | (0, 0.2596) | 1.2788 | 0.0142 | 0.0597 |
| Nodule | M+R+ | 54.4899 | 134.9300 | 0.2877 | (0.1499, 0.4155) | 11.852 | 0.7845 | 0.6228 |
| | M-R+ | 124.4200 | 133.2200 | 0.4829 | (0.3404, 0.6060) | 14.095 | 0.9382 | 0.7913 |

**NOTE**: Broad sense hertability ($H^2$) was calculated by dividing the genetic contribution to plant phenotype ($V_g$) by the total variation in the phenotype ($V_g$+$V_e$). $V_g$ was the variation among genotypes in our model and $V_e$ (variation in phenotype due to the environment) was the residual variance in our models. 95% CIs around heritabilities were calculated using a randomization with 10,000 draws from a bivariate normal distribution. $\bar{X}$ and $\bar{X}$ SE represent the mean and standard error of trait values from raw data and CVG represents the Coefficient of Variation (calculated by diving the square root of $V_g$ by $\bar{X}$). 'Trans' denotes square root plus one transformed response data.

**Supplementary Table S5: Effects of a third-party mutualist (*i.e.* mycorrhizal fungi) on fitness alignment between rhizobia and their host**

***(a) Effect of mycorrhizal fungi on the relationship between rhizobia and plant fitness***

| **Factor** | **DF** | **F** | **P** |
|---|---|---|---|
| mycorrhizal fungi (myco) | 1 | 2.58 | 0.1089 |
| nodule number (nodule) | 1 | 28.49 | **1.69E-07** |
| myco × nodule | 1 | 4.94 | **0.0269** |
| Residuals | 355 | | |

**NOTE**: "mycorrhizal fungi" ("myco") indicates treatment manipulating presence/absence of mycorrhizal fungi, "nodule number" is the mean number of nodules for a given genotype in the relevant microbial environment. Significant p-values are in bold.

***(b) Effect within each microbial environment***

| | **DF** | **F** | **P** |
|---|---|---|---|
| **M+R+** | 1, 182 | 21.45 | 6.90E-06 |
| **M-R+** | 1, 173 | 6.90 | 0.0094 |

**NOTE**: This provides information about the regressions between nodule number and pod number in each microbial environment separately.

**Supplementary Table S6: Test of interactive effects of multiple microbial mutualists on selection on host traits**

| Factor | DF | F | P |
|---|---|---|---|
| **mycorrhizal fungi (myco)** | **1** | **11.11** | **0.0009** |
| rhizobia (rhizo) | 1 | 2.52 | 0.1132 |
| **Branching** | **1** | **60.65** | **2.62E-14** |
| **root:shoot** | **1** | **3.06** | **0.0809** |
| **myco × rhizo** | **1** | **10.69** | **0.0011** |
| **rhizo × branching** | **1** | **44.43** | **5.55E-11** |
| **myco × branching** | **1** | **5.92** | **0.0152** |
| **rhizo × root:shoot** | **1** | **3.43** | **0.0646** |
| myco × root:shoot | 1 | 0.01 | 0.9399 |
| **myco × rhizo × branching** | **1** | **5.71** | **0.0171** |
| myco × rhizo × root:shoot | 1 | 0.02 | 0.8826 |
| Residuals | 664 | | |

**NOTE**: "mycorrhizal fungi" ("myco") indicates treatment manipulating presence/absence of mycorrhizal fungi, "rhizobia" ("rhizo") indicates treatment manipulating presence/absence of rhizobia, "branching" is the number of branches the plant has (calculated mean value for the genotype), and "root:shoot" is the ratio of below to aboveground plant biomass (calculated mean value for the genotype). Significant (and marginally significant) p-values are in bold.

**Supplementary Table S7: Test of interactive effects of multiple microbial mutualists on selection on host traits (using plant fitness relativized within treatment)**

| Factor | DF | F | P |
|---|---|---|---|
| mycorrhizal fungi (myco) | 1 | 0.04 | 0.8344 |
| rhizobia (rhizo) | 1 | 0.09 | 0.7698 |
| **branching** | **1** | **100.83** | **<2.2E-16** |
| root:shoot | 1 | 0.07 | 0.7967 |
| myco × rhizo | 1 | 0.01 | 0.9365 |
| **rhizo × branching** | **1** | **17.29** | **3.63E-5** |
| **myco × branching** | **1** | **4.50** | **0.0343** |
| rhizo × root:shoot | 1 | 1.36 | 0.2435 |
| myco × root:shoot | 1 | 0.11 | 0.7435 |
| **myco × rhizo × branching** | **1** | **6.32** | **0.0122** |
| myco × rhizo × root:shoot | 1 | 0.09 | 0.7586 |
| Residuals | 664 | | |

**NOTE**: "mycorrhizal fungi" ("myco") indicates treatment manipulating presence/absence of mycorrhizal fungi, "rhizobia" ("rhizo") indicates treatment manipulating presence/absence of rhizobia, "branching" is the number of branches the plant has (calculated mean value for the genotype), and "root:shoot" is the ratio of below to aboveground plant biomass (calculated mean value for the genotype). Relative fitness was calculated for each treatment separately, and traits were mean centered with stdev=1 within each treatment. Significant (and marginally significant) p-values are in bold.

**Supplementary Table S8**: **Univariate analyses to determine selection gradients for microbial effects on plant traits.**

| Treatment | Trait | S | S.E. | P-value | 95% CIs | Trait Mean | Trait S.D. | Mean Fitness |
|---|---|---|---|---|---|---|---|---|
| M+R+ | **Branching** | **0.515** | **0.039** | **2.00E-16** | **( 0.34, 0.73 )** | **7.25** | **5.58** | 2.27 |
| | **Root:Shoot** | **-1.576** | **0.715** | **0.0290** | **(-3.17, -0.72 )** | **0.46** | **0.47** | |
| M-R+ | **Branching** | **0.272** | **0.034** | **9.82E-14** | **( 0.18, 0.36 )** | **7.06** | **5.38** | 2.16 |
| | **Root:Shoot** | **-1.042** | **0.433** | **0.0173** | **(-1.93, -0.48 )** | **0.52** | **0.52** | |
| M+R- | **Branching** | **0.033** | **0.013** | **0.0098** | **( 0.01, 0.06 )** | **2.78** | **1.69** | 0.14 |
| | Root:Shoot | 0.015 | 0.032 | 0.6302 | (-0.04, 0.08 ) | 0.84 | 0.70 | |
| M- R- | **Branching** | **0.032** | **0.012** | **0.0113** | **( 0.01, 0.06 )** | **2.62** | **1.93** | 0.13 |
| | Root:Shoot | -0.016 | 0.038 | 0.6770 | (-0.06, 0.03) | 0.71 | 0.66 | |

**NOTE:** In each of the four microbial environments, selection was calculated (‘S’) and its significance (denoted under ‘P-value’) was determined using univariate linear regression analyses within each of the microbial treatments for each trait. Bias corrected 95% CIs around each selection gradient were calculated using 10,000 bootstraps. Reported trait means and standard deviations for each treatment were calculated across the genotype means within that treatment. For treatments, M and R indicate mycorrhizal fungi and rhizobia, and + and – indicate presence or absence of that microbe. Significant selection on a trait is indicated by bold text.
